# Dorsal Raphe to Basolateral Amygdala Corticotropin-Releasing Factor Circuit Regulates Cocaine-Memory Reconsolidation

**DOI:** 10.1101/2024.02.10.579725

**Authors:** Jobe L. Ritchie, Shuyi Qi, David A. Soto, Sydney E. Swatzell, Hope I. Grenz, Avery Y. Pruitt, Lilia M. Artimenia, Spencer K. Cooke, Craig W. Berridge, Rita A. Fuchs

## Abstract

Environmental stimuli elicit drug craving and relapse in cocaine users by triggering the retrieval of strong cocaine-related contextual memories. Retrieval can also destabilize drug memories, requiring reconsolidation, a protein synthesis-dependent storage process, to maintain memory strength. Corticotropin-releasing factor (CRF) signaling in the basolateral amygdala (BLA) is necessary for cocaine-memory reconsolidation. We have hypothesized that a critical source of CRF in the BLA is the dorsal raphe nucleus (DR) based on its neurochemistry, anatomical connectivity, and requisite involvement in cocaine-memory reconsolidation. To test this hypothesis, male and female Sprague-Dawley rats received adeno-associated viruses to express Gi-coupled designer receptors exclusively activated by designer drugs (DREADDs) selectively in CRF neurons of the DR and injection cannulae directed at the BLA. The rats were trained to self-administer cocaine in a distinct environmental context then received extinction training in a different context. They were then briefly re-exposed to the cocaine-predictive context to destabilize (reactivate) cocaine memories. Intra-BLA infusions of the DREADD agonist deschloroclozapine (DCZ; 0.1 mM, 0.5 μL/hemisphere) after memory reactivation attenuated cocaine-memory strength, relative to vehicle infusion. This was indicated by a selective, DCZ-induced and memory reactivation-dependent decrease in drug-seeking behavior in the cocaine-predictive context in DREADD-expressing males and females at test compared to respective controls. Notably, BLA-projecting DR CRF neurons that exhibited increased c-Fos expression during memory reconsolidation co-expressed glutamatergic and serotonergic neuronal markers. Together, these findings suggest that the DR_CRF_ → BLA circuit is engaged to maintain cocaine-memory strength after memory destabilization, and this phenomenon may be mediated by DR CRF, glutamate, and/or serotonin release in the BLA.

## INTRODUCTION

Environmental stimuli can become occasion setters that predict access to drugs of abuse upon the formation of enduring context-response-drug associative memories [1,2]. The retrieval of such drug memories can elicit drug craving and relapse in individuals with substance use disorder (**SUD**), but it can also temporarily destabilize those memories [3]. Labile drug memories must be reconsolidated into long-term memory stores to endure [3], and disruption of labile drug memories or interference with their reconsolidation curtails stimulus control over drug-seeking behavior in animal models of drug relapse [4,5] and drug-cue reactivity in substance users [6-9]. Thus, therapeutic strategies that target labile drug memories or reconsolidation may be promising for relapse prevention [10], but refinement of such approaches requires deeper understanding of underlying neurobiological mechanisms.

The basolateral amygdala complex (**BLA**) is a site of protein synthesis for cocaine-memory reconsolidation [11]. Cocaine-memory reconsolidation requires the activation of intracellular signaling molecules, including protein kinase A [12,13] and extracellular signal-regulated kinase [14], and the expression of immediate early-genes (**IEGs**), including zinc finger 268 (**Zif268**) [4,15], in the BLA. Upstream from these signaling events, corticotropin releasing factor (**CRF**) signaling in the BLA is necessary and sufficient for cocaine-memory reconsolidation. In support of this, intra-BLA CRF receptor type 1 (**CRFR1**) antagonist administration attenuates cocaine-memory strength, independent of sex, and exogenous CRF administration enhances memory strength in female rats [16]. However, the critical source of BLA CRF during cocaine-memory reconsolidation has not been investigated.

The dorsal raphe (**DR**) is a potential source of CRF to the BLA based on its neurochemistry and involvement in cocaine-memory reconsolidation. DR neuronal populations can release several neurotransmitters, including glutamate, serotonin, dopamine, and CRF, that have been implicated in memory reconsolidation [16-19] and alter their firing rate upon the presentation of reward-predictive stimuli [20]. We have demonstrated that DR engagement is critical for cocaine-memory reconsolidation, as GABA agonist-induced inhibition of DR neuronal signaling after memory reactivation weakens cocaine memories and diminishes subsequent context-induced cocaine-seeking behavior [21]. However, the role of DR projections, and specifically DR *CRF* inputs to the BLA (**DR**_**CRF**_**→BLA**), in cocaine-memory reconsolidation is not known.

Here, we tested the hypothesis that DR_CRF_→BLA circuit function is necessary for cocaine-memory reconsolidation, using cell- and circuit-specific chemogenetic inhibition in rats. To this end, we expressed inhibitory designer receptors exclusively activated by designer drugs (**DREADDs**) in DR CRF neurons and aimed injection cannulae at the BLA. The rats underwent cocaine self-administration training in a salient context, to establish context-response-cocaine memories, and then extinction training in a different context. They were then briefly re-exposed to the cocaine-predictive context to destabilize (*reactivate*) cocaine memories. Rats received intra-BLA deschloroclozapine (**DCZ**; DREADD agonist) treatment immediately after cocaine-memory reactivation (while the memories were labile) or 6 hours later (after the memories were reconsolidated) [22]. Moreover, we determined the cell types of BLA-projecting DR CRF neurons that exhibited activation, as indicated by increased c-Fos expression, during cocaine-memory reconsolidation.

## METHODS

### Additional methodological details are provided in Supplemental Materials and Methods

#### Subjects

Male (*n* = 47; body weight: 275–325 g) and female (*n* = 45; body weight: 220–250 g) Sprague-Dawley rats were housed individually on a reversed light cycle (lights off at 6:00 a.m.). Rats received unlimited access to water and 20–25 g of standard rat chow per day. Animal housing and care protocols followed the *Guide for the Care and Use of Laboratory Animals* [23] and were approved by the Washington State University Institutional Animal Care and Use Committee.

#### Surgery

Rats were anesthetized using ketamine and xylazine (100.0 and 5.0 mg/kg, i.p., respectively) at least 24 hours after a food-training session (see *Supplement*). Back-mounted intravenous catheters were implanted into the right jugular vein as described previously [24]. Rats in Experiments 1-2 then received a unilateral virus-cocktail infusion into the median DR (30° angle to target the midline) and stainless-steel guide cannula implants (P1 Technologies, Roanoke, VA) aimed bilaterally 2 mm dorsal to the BLA. The virus cocktail (200 nL) contained, in 2:1 ratio, AAV8-rCRHp-iCre (1.2 x 10^13^ GC/mL, Vector Biolabs, Malvern, PA) plus either AAV8-hSyn-DIO-hM4Di-mCherry (2.2 x 10^13^ GC/mL, Addgene, Watertown, MA) or AAV8-hSyn-DIO-mCherry (Control virus; 2.3 x 10^13^ GC/ML, Addgene). Rats in Experiment 3 received a unilateral infusion of AAV8-rCRHp-iCre (200 nL; 1.2 x 10^13^ GC/mL, Vector Biolabs) into the DR and bilateral infusions of AAVretro-hSyn-DIO-EGFP (200 nL over 2 min; 1.3 x 10^13^ GC/mL, Addgene) into the BLA. Post-operative analgesic treatment was provided during the first 48 hours of the five-day post-surgical recovery period (see *Supplement*).

#### Drug self-administration and extinction training

Drug self-administration training was conducted six days per week in operant-conditioning chambers (Coulbourn, Allentown, PA) that contained salient auditory, visual, tactile, and olfactory stimuli. These two-hour sessions took place during the rats’ dark phase. Active-lever responses resulted in cocaine reinforcement (0.5 mg/kg per 50-μL infusion, delivered i.v. over 2 seconds; generously provided by the NIDA Drug Supply Program, Research Triangle Park, NC) under a fixed ratio 1 schedule with a 20-second timeout period after each infusion. Inactive-lever responses were not reinforced at any time. Training continued until the rat received at least 10 sessions during which they obtained ≥ 10 infusions/session. Rats then received seven daily 2-hour extinction training sessions in a distinctly different context, where lever responses were not reinforced. Rats in Experiments 1-2 were acclimated to the intracranial infusion procedure immediately after extinction session 4 (see *Supplement*).

#### Memory reactivation and DREADD agonist treatment

Rats were placed into the cocaine-predictive context for 15 min to elicit the retrieval and destabilization of contextual cocaine memories (memory reactivation) or remained in their home cages (no-memory reactivation). Group assignment was balanced based on mean active-lever responding on the last three cocaine self-administration days. The 15-minute session length is sufficient to destabilize cocaine memories without eliciting overt behavioral extinction [11,21]. The rats were connected to the infusion lines, but lever responses were not reinforced. Rats received intra-BLA infusions of saline vehicle (VEH; 0.5 μl/hemisphere) or DCZ (0.1 mM; 0.5 μl/hemisphere; HB9126, Hello Bio, Princeton, NJ) immediately or six hours later (Experiments 1-2), or they were euthanized without intra-BLA treatment (Experiment 3) (see *Supplement* for intracranial infusion protocol).

#### Tests of memory strength

One day after intra-BLA treatment, rats in Experiments 1-2 received daily two-hour extinction training sessions until their active-lever responding declined to ≤ 25 on two consecutive days. They then received a two-hour test session in the cocaine-predictive context. Active-lever responses upon first re-exposure to the extinction and cocaine-predictive contexts served as indices of extinction and cocaine-memory strength, respectively. Lever responding was not reinforced during testing.

#### Brain histology

Rats received an overdose of ketamine and xylazine and then transcardial perfusion as described previously [21] (see *Supplement*). Brains were extracted, cryoprotected, and stored at −80 °C. The brains were sectioned on a cryostat (Leica Biosystems, Buffalo Grove, Illinois, USA). Thirty-μm brain sections were stained with DAPI (1:1000; D1306, Invitrogen) and mounted on glass slides. Injection-cannula placements and virus spread were assessed using an epifluorescence microscope (Leica CTR 6500, Wetzler, Germany).

#### *In situ* hybridization

The CRF antibody was validated using three digoxigenin-labeled 20-nucleotide RNA probes (MicroSynth, Balgach, Switzerland) directed against segments of rat CRF mRNA (NCBI GenBank reference sequence: NM_031019.2). The probes were hybridized with CRF mRNA as described previously [25], and bound probes were visualized using immunolabeling (see *Supplement* for details).

#### Immunohistochemistry

Standard protocols were used to fluorescently label targets in free-floating brain slices ([21,25]; see *Supplement* for details). Briefly, brain slices were placed in blocking solution for 1 hour then incubated with an appropriate primary antibody for 20 or 72 h at 4°C (for CRF protein) or overnight at room temperature (for IEGs and cell-type markers) (antibody information is provided in ***Table S1***). They were washed in PBS then incubated with appropriate secondary antibodies for 2 hours. Multiple targets were immunolabeled sequentially, across days, to avoid antibody interactions.

#### Image analysis

Brain tissue was imaged using epifluorescence (10x dry; Leica CTR 6500) or confocal (40x or 63x oil immersion; Leica SP8 or Nikon A1R+, Melville, NY) microscopy (see *Supplement*). Single-labeled BLA cell bodies were quantified on four images per subject using a custom macro in ImageJ. Multi-labeled DR cell bodies were quantified on three z-stacks (28-μm span, 1-μm steps) per DR subregion per subject and counted in ImageJ by two independent observers blinded to the subjects’ treatment condition (inter-observer reliability in Experiment 3, R^2^ = 0.96). All data were converted to density values and averaged across replicates.

### Data analysis

Data from rats that failed to reach behavioral training criteria or had misplaced guide cannulae or misplaced/insufficient virus expression (*n* = 17, 12, and 23 respectively) were excluded from analysis. Lever responses, drug infusions, response latency (i.e., time to the third active-lever response [26]), and cell density were analyzed using mixed-factorial analyses of variance (ANOVA) – with treatment group [(Gi) VEH, (Gi) DCZ, or (Ctrl) DCZ], memory-reactivation condition (no-memory reactivation, memory reactivation), and sex as between-subjects factors and testing context (extinction, cocaine-predictive) and time (sessions, 20-min bins) as within-subject factors – or *t*-tests, where appropriate. Significant interaction and time main effects were probed using Bonferroni’s *post hoc* tests. Alpha was set at 0.05.

## RESULTS

### Behavioral history

There were no pre-existing differences between the groups in Experiments 1-3 in lever responding or drug intake during self-administration training, extinction training, or the memory-reactivation session (**Figs. 2C, 3C, 4B; see** statistics in ***Table S2***). Furthermore, DCZ treatment did not alter the number of session rats needed to reach the extinction criterion (mean ± SEM = 2.15 ± 0.08 sessions).

**Figure 1.**
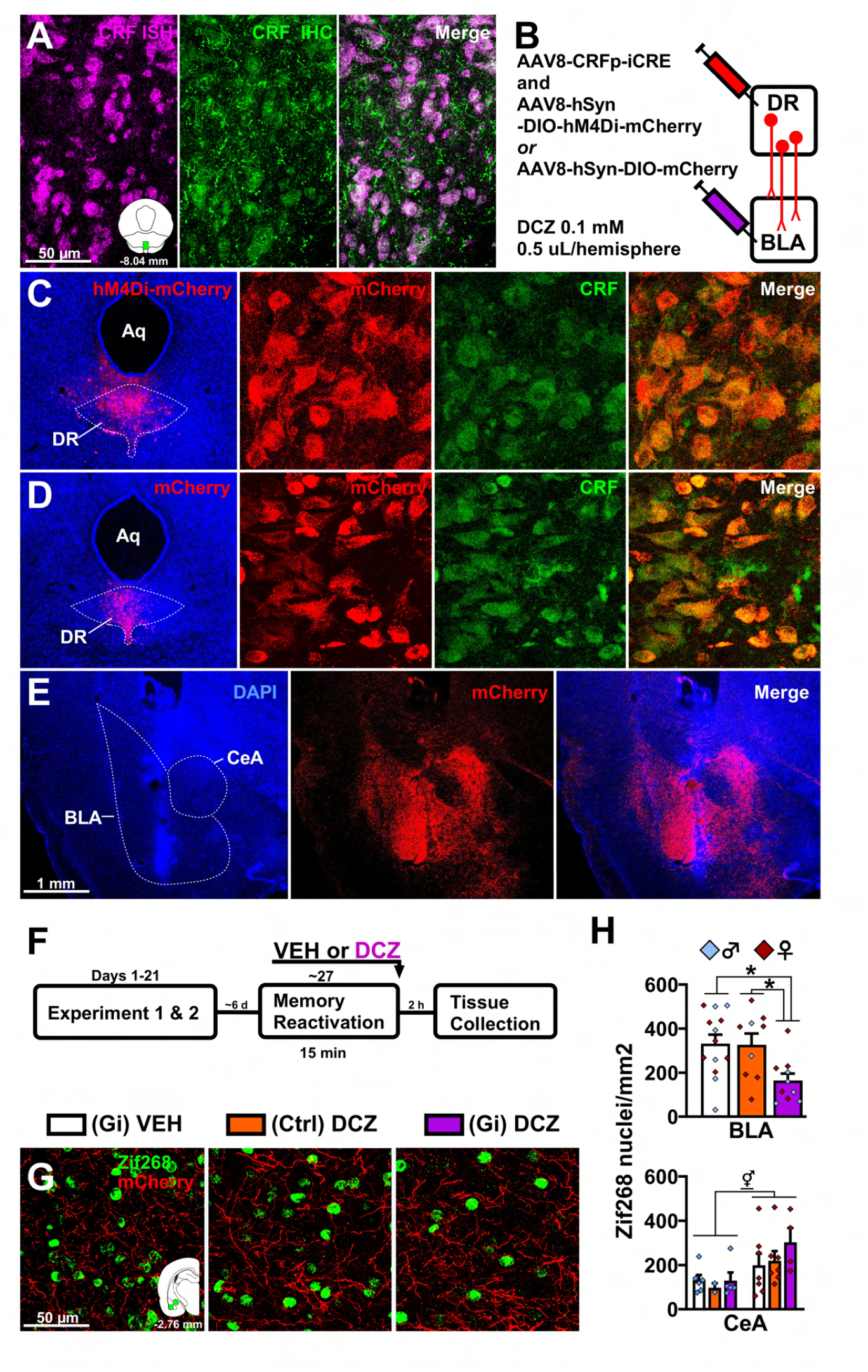
Validation study results. **(A)** Corticotropin-releasing factor (**CRF**) antibody specificity as indicated by colocalization between CRF mRNA, visualized using fluorescence *in situ* hybridization (**CRF ISH**), and CRF protein, labeled using immunohistochemistry (**CRF IHC**), in 97.1 ± 0.9 % of the DR cell bodies examined. **(B)** Schematic of the cell- and circuit-specific chemogenetic approach. Rats received an infusion of a cocktail of viruses (200 nL) into the midline dorsal raphe (**DR**) to express the hM4Di fused with mCherry (**Gi**) or mCherry alone (Control; **Ctrl**) Cre-dependently, under the CRF promoter. Guide cannulae were implanted into the basolateral amygdala (**BLA**) to permit bilateral infusions of deschloroclozapine (**DCZ**) in the terminal region of the target circuit. **(C)** Representative hM4Di-mCherry expression in the DR and colocalization of hM4Di-mCherry expression with CRF-immunoreactivity seen in 96.9 ± 0.6 % of DR mCherry-expressing neurons examined. **(D)** Representative mCherry expression in the DR and colocalization of mCherry expression with CRF-immunoreactivity. **(E)** Representative mCherry-expressing DR CRF axon terminals in the BLA and adjacent central amygdaloid nucleus (**CeA**) and injection cannula tract in the BLA. **(F)** Experimental timeline for DREADD efficacy and anatomical selectivity assessment. A subset of rats received a memory-reactivation session ∼6 days after their last test session in Experiment 1 or 2. Memory reactivation was followed immediately by intra-BLA VEH or DCZ treatment [(Gi) VEH: *n* = 6 males, 7 females; (Ctrl) DCZ: *n* = 2 males, 7 females; (Gi) DCZ: *n* = 5 males, 4 females)]. Brain tissue was collected 2 hours later to assess Zif268 expression in the BLA and adjacent CeA. **(G)** Representative images of Zif268-immunoreactive nuclei and mCherry expression in the BLA. **(H)** Density of Zif268 expression (mean number of nuclei/mm^2^ ± SEM) in the BLA (upper panel) and CeA (lower panel). ***Symbols***: ANOVA *treatment group main effect or t-test; ^⚥^sex main effect (underlined). All *p*s < 0.05. ***Abbreviations*:** Aq, cerebral aqueduct.

**Figure 2.**
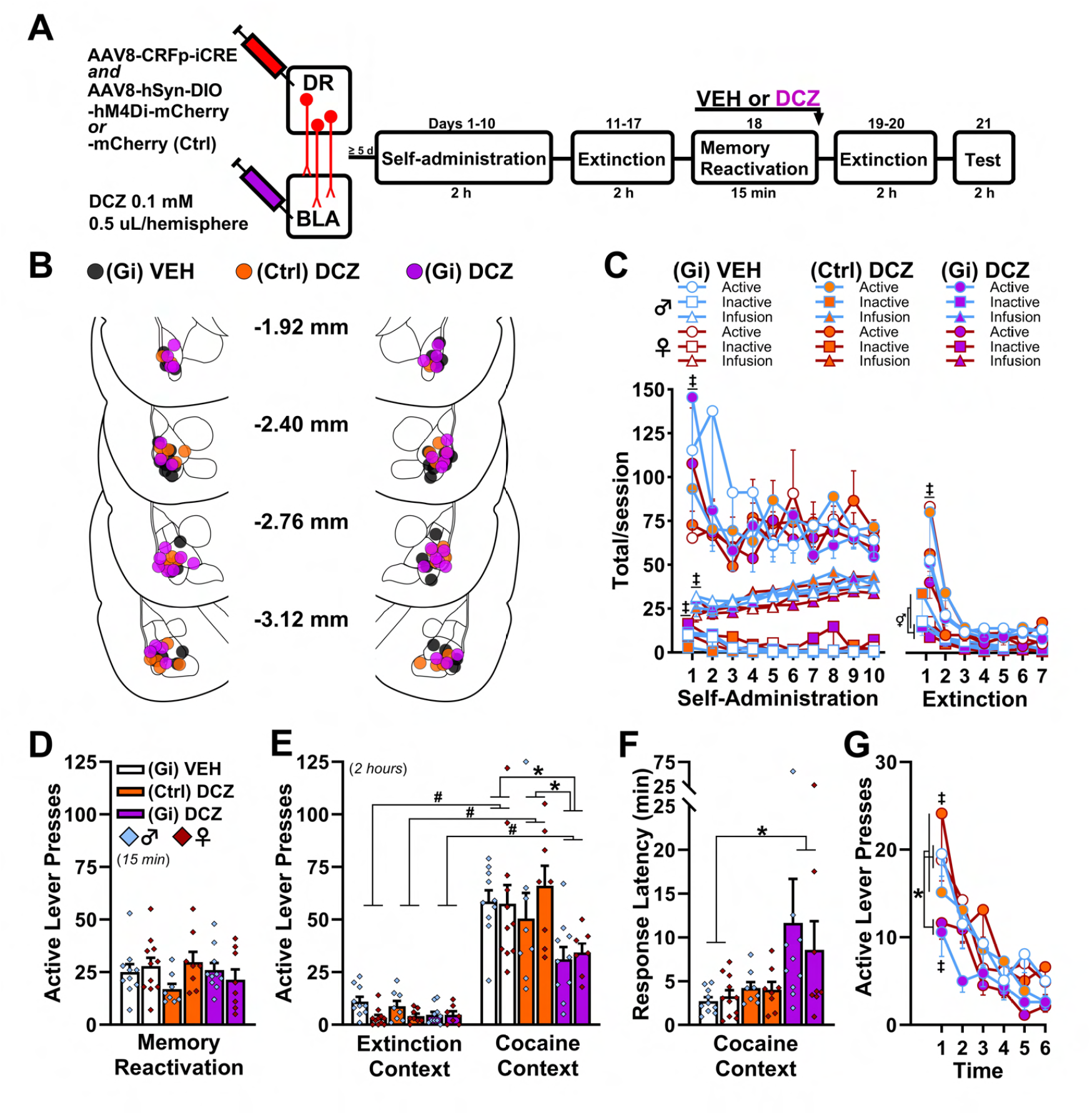
Chemogenetic DR_CRF_ → BLA circuit inhibition prior to memory reconsolidation weakens cocaine-memory strength. **(A)** Timeline of Experiment 1. The Gi-coupled DREADD hM4Di fused with mCherry (**Gi**) or mCherry alone (control; **Ctrl**) was expressed in dorsal raphe (**DR**) corticotropin-releasing factor (**CRF**) neurons and guide cannulae were directed at the basolateral amygdala (**BLA**). Rats then received cocaine self-administration training in a salient environmental context and extinction training in a different context. Next, they received 15-minute re-exposure to the cocaine-predictive context (i.e., memory reactivation) followed by bilateral intra-BLA microinjections of vehicle (**VEH**) [(Gi) VEH; *n* = 10 males, 11 females)] or the DREADD agonist deschloroclozapine (**DCZ**) [(Ctrl) DCZ: *n* = 8 males, 8 females; (Gi) DCZ: *n* = 10 males, 8 females)]. Non-reinforced lever responding in the extinction and cocaine-predictive contexts was tested 24 and 72 hours later, respectively. **(B)** Injection cannula placements in the BLA. The numbers between the schematics indicate AP distance from Bregma in millimeters. **(C)** Cocaine infusions and active- and inactive-lever responses (mean/2 h ± SEM) during cocaine self-administration training (last 10 sessions) and extinction training. **(D)** Active-lever responses during the memory-reactivation session (mean/15 minutes ± SEM) immediately prior to intra-BLA treatment. **(E)** Active-lever responses (mean/2 hours ± SEM) upon first re-exposure to the extinction and cocaine-predictive contexts after intra-BLA treatment. **(F)** Active-lever response latency (mean ± SEM) in the cocaine-predictive context at test. **(G)** Time course of active-lever responses (mean/20 min ± SEM) in the cocaine-predictive context at test. ***Symbols***: ANOVA ^‡^time main (underlined) or simple main effect (see details for pairwise comparisons in *Results*); ^⚥^sex simple main effect; ^#^context simple main effect; *treatment group simple main effect. All *p*s ≤ 0.05.

**Figure 3.**
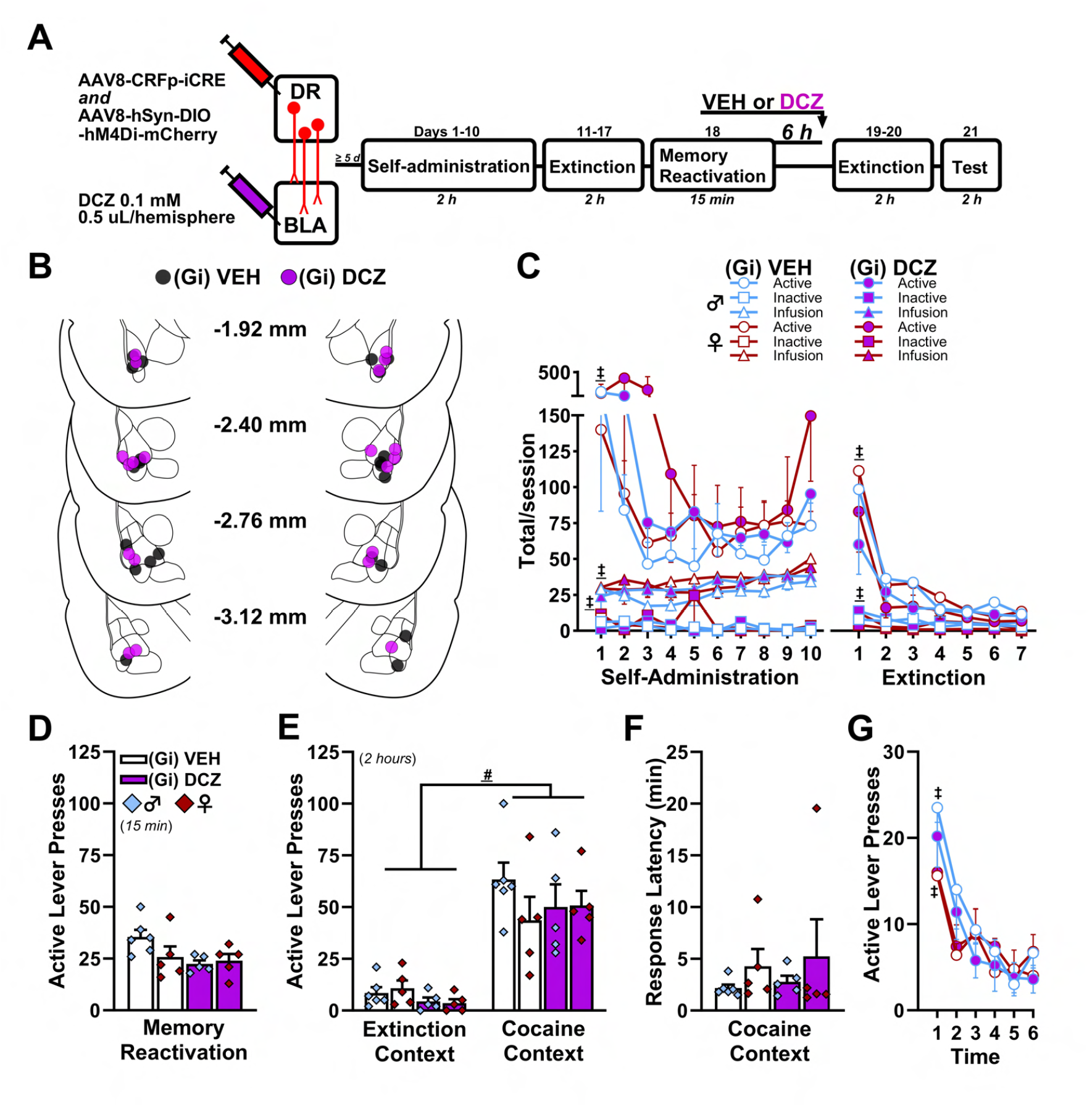
Chemogenetic inhibition of the DR_CRF_ → BLA circuit after memory reconsolidation does not alter cocaine-memory strength. **(A)** Timeline of Experiment 2. The Gi-coupled DREADD hM4Di fused with mCherry (**Gi**) was expressed in dorsal raphe (**DR**) corticotropin-releasing factor (**CRF**) neurons and guide cannulae were directed at the basolateral amygdala (**BLA**). Rats then received cocaine self-administration training in a distinct context and extinction training in a different context. Next, they received 15-minute re-exposure to the cocaine-predictive context (i.e., memory reactivation). Six hours later, the rats received bilateral intra-BLA injections of vehicle (**VEH**) [(Gi) VEH; *n* = 6 males, 5 females)] or the DREADD agonist deschloroclozapine (**DCZ**) [(Gi) DCZ; *n* = 5 males, 5 females)]. Non-reinforced lever responding in the extinction and cocaine-predictive contexts was tested 24 and 72 hours later, respectively. **(B)** Injection cannula placements in the BLA. The numbers between the schematics indicate AP distance from Bregma in millimeters. **(C)** Cocaine infusions and active- and inactive-lever responses (mean/2 h ± SEM) during cocaine self-administration training (last 10 sessions) and extinction training. **(D)** Active-lever responses during the memory-reactivation session (mean/15 minutes ± SEM) 6 hours prior to intra-BLA treatment. **(E)** Active-lever responses (mean/2 hours ± SEM) upon first re-exposure to the extinction and cocaine-predictive contexts after delayed intra-BLA treatment. **(F)** Active-lever response latency (mean ± SEM) in the cocaine-predictive context at test. **(G)** Time course of active-lever responses (mean/20 min ± SEM) in the cocaine-predictive context at test. ***Symbols***: ^‡^time main (underlined) or simple main effect (see details for pairwise comparisons in Results); ^#^context main effect; All *p*s < 0.05.

**Figure 4.**
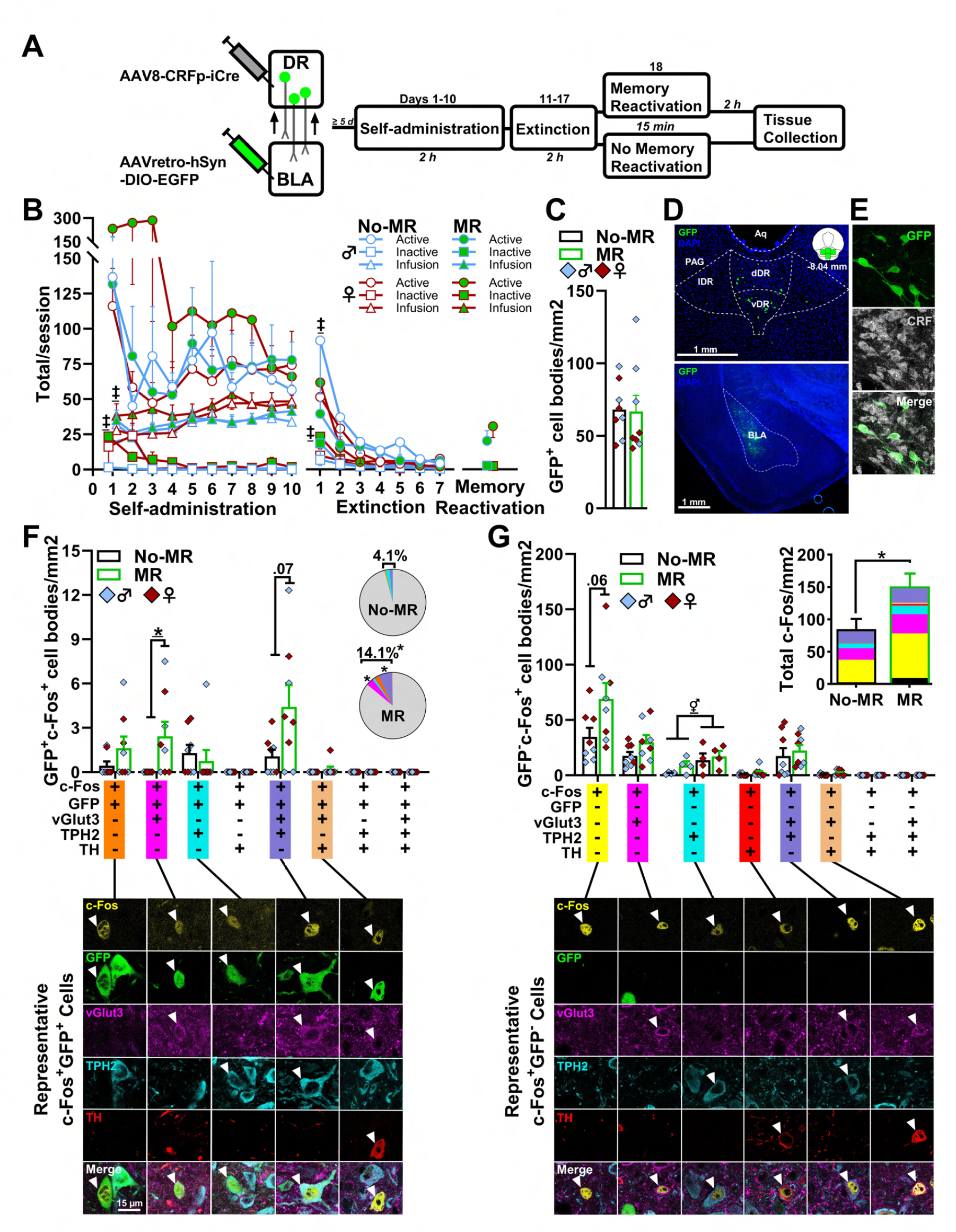
BLA-projecting DR CRF neurons activated during cocaine-memory reconsolidation co-express vGlut3 and TPH2. **(A)** Timeline of Experiment 3. GFP was expressed in basolateral amygdala (**BLA**) -projecting corticotropin-releasing factor (**CRF**) cell bodies in the dorsal raphe (**DR**) using a combinatorial viral approach. Rats then received cocaine self-administration training in a distinct context and extinction in a different context. Next, they received 15-minute re-exposure to the cocaine-predictive context to reactive cocaine memories (**MR**; *n* = 4 males, 4 females) or remained in their home cages (**No-MR;** *n* = 4 males, 4 females). Brain tissue was collected 2 hours later to capture c-Fos expression approximately 30 minutes into memory reconsolidation. **(B)** Cocaine infusions and active- and inactive-lever responses (mean/2 h ± SEM) during cocaine self-administration training (last 10 sessions), extinction training sessions, and memory reactivation. **(C)** GFP-expressing (BLA-projecting CRF) cell body density in the DR (mean cell bodies/mm^2^ ± SEM) in the No-MR and MR groups. **(D)** Representative image of GFP expression in BLA-projecting DR neurons and in their axon terminals within the BLA of the same subject. **(E)** Colocalization between GFP-expression and CRF-immunoreactivity seen in 98 ± 0.2% of DR neurons examined. **(F, upper panel)** Density of c-Fos^+^ GFP-expressing (GFP^+^) neuronal populations in the DR, following No-MR and MR (mean cell bodies/mm^2^ ± SEM). Data are shown collapsed across sex where sex differences were not observed. (**F, Inset**) Proportion of c-Fos^+^ GFP-expressing neurons following No-MR and MR. (**F, lower panel**) Representative images of c-Fos^+^ GFP-expressing cell types observed in the DR. **(G, upper panel**) Density of c-Fos^+^ GFP-non-expressing (GFP^-^) cell types following No-MR and MR (mean cell bodies/mm^2^ ± SEM). Data are shown collapsed across sex where sex differences were not observed. (**G, Inset**) Total c-Fos expression (mean GFP^+^ plus GFP^-^ cell bodies/mm^2^ ± SEM) following No-MR and MR. Black segment represents cFos^+^ GFP^+^ neurons collapsed across cell type. (**G, lower panel**) Representative images of c-Fos^+^ GFP-non-expressing cell types observed in the DR. ***Symbols***: ^‡^time main effect; *treatment group simple main effect (underlined) or *t*-test, No-MR vs MR; ^⚥^sex main effect; All *p* < 0.05. ***Abbreviations*: Aq**, cerebral aqueduct; **dDR**, dorsal subregion of dorsal raphe; **lDR**, lateral subregion of dorsal raphe; **vDR**, ventral subregion of dorsal raphe; **PAG**, periaqueductal gray.

### Validation studies

CRF immunoreactivity colocalized with CRF mRNA expression in 97.1 ± 0.9 % of DR cell bodies (**Fig. 1A**), confirming antibody selectivity. Similarly, CRFp-iCre-dependent mCherry (+/- hM4Di) expression colocalized with CRF immunoreactivity in 96.9 ± 0.6 % of DR cell bodies (**Fig. 1C-D**; [27]) and was observed in DR terminals within the BLA (**Fig. 1E**), indicating cell-specific transgene expression. To evaluate the functionality and anatomical specificity of chemogenic manipulations (**Fig. 1F-H**), some rats from Experiments 1-2 (*n* = 2 - 7/sex/group) received a 15-minute memory-reactivation session followed immediately by intra-BLA VEH or DCZ treatment. BLA brain tissue was collected 2 hours later to assess Zif268 expression[28], which is necessary in the BLA for cocaine-cue memory reconsolidation [4,15]. DCZ in the h4MDi-expressing group [**(Gi) DCZ)**] reduced BLA Zif268 expression independent of sex (**Fig. 1G-H, upper panel**; 2 x 3 ANOVA; treatment main effect only, F_(2,25)_ = 4.30, p = 0.03; all other *F*s ≤ 0.31, *p* ≥ 0.59), relative to VEH in h4MDi-expressing group [**(Gi) VEH**; Bonferroni’s test, *p* = 0.04)] and DCZ in mCherry-expressing controls [**(Ctrl) DCZ**; *t*_(16)_ = 2.76, *p* = 0.02)]. Conversely, intra-BLA DCZ did not alter Zif268 expression in the central amygdaloid nucleus (**CeA**), where females exhibited greater Zif268 expression than males independent of treatment condition (**Fig. 1H, lower panel;** sex main effect only, *F*_(1,25)_ = 7.91, *p* = 0.009; all other *F*s ≤ 0.73, *p* ≥ 0.49).

### DR_CRF_ → BLA circuit inhibition weakens labile cocaine memories

Experiment 1 examined whether DR_CRF_ → BLA circuit function was necessary for cocaine-memory reconsolidation (**Fig. 2;** see ***Fig. S1*** for inactive lever results). Rats received intra-BLA VEH or DCZ treatment immediately after memory reactivation, when cocaine memories were expected to be labile [4]. At test, the cocaine-predictive context elicited more active-lever responding than the extinction context, independent of sex (**Fig. 2E**; 2 x 3 x 2 ANOVA; context x treatment interaction, *F*_(2,49)_ = 5.29, *p* = 0.01, Bonferroni’s tests, *p*s < 0.05; context main effect, *F*_(1,49)_ = 162.30, *p* < 0.001; treatment main effect, *F*_(2,49)_ = 7.83, *p* = 0.001; all other *F*s ≤ 2.08, *p*s ≥ 0.16). Collapsed across sex, DCZ reduced active-lever responding in the h4MDi-expressing group selectively in the cocaine-predictive context, relative to (Gi) VEH or (Ctrl) DCZ control groups (Bonferroni’s tests, *p*s < 0.05). DCZ also augmented active-lever response latency in the cocaine-predictive context in the hM4Di-expressing group compared to the (Gi) VEH group, independent of sex (**Fig. 2F**; 2 x 3 ANOVA; treatment main effect, *F*_(2,49)_ = 4.39, Bonferroni’s test, *p* = 0.02, *p* = 0.02; all other *F*s ≤ 0.28, *p* ≥ 0.76), while the (Ctrl) DCZ group did not differ from either group. Time-course analysis indicated that active-lever responding was highest during the first 20 minutes in the cocaine-predictive context independent of sex or treatment group (**Fig. 2G**; 2 x 3 x 6 ANOVA; time main effect, *F*_(5,245)_ = 38.50, *p* < 0.001, bin 1 > 2-6, Bonferroni’s tests, *p*s < 0.05; treatment main effect, *F*_(2,49)_ = 6.79, *p* = 0.002; all other *F*s ≤ 1.49, *p*s ≥ 0.14). Collapsed across sex and time, DCZ reduced responding in the hM4Di-expressing group relative to (Gi) VEH and (Ctr) DCZ control groups (Bonferroni’s tests, *p*s < 0.05).

### DR_CRF_ → BLA circuit inhibition does not disrupt reconsolidated cocaine memories

Experiment 2 assessed whether DR_CRF_ → BLA circuit inhibition would alter the strength of already reconsolidated cocaine memories (**Fig. 3**). Rats received intra-BLA VEH or DCZ treatment six hours after memory reactivation, when cocaine memories were expected to be stable [22]. At test, the cocaine-predictive context elicited more active-lever responding than the extinction context, independent of sex or delayed DCZ treatment (**Fig. 3E**; 2 x 2 x 2 ANOVA; context main effect only, *F*_(1,17)_ = 79.47, *p* < 0.001; all other *F*s ≤ 1.36, *p*s ≥ 0.26). Neither sex nor delayed DCZ treatment altered response latency in the cocaine-predictive context (**Fig. 3F**; 2 x 2 ANOVA, *F*s ≤ 1.42, *p*s ≥ 0.25). Time-course analysis indicated that active-lever responding peaked during the first 20 minutes in the cocaine-predictive context independent of sex or delayed DCZ treatment (**Fig. 3G**; 2 x 2 x 6 ANOVA; sex x time interaction, *F*_(5,85)_ = 2.46, *p* = 0.04; time main effect, *F*_(5,85)_ = 24.01, *p* < 0.001; all other *F*s ≤ 1.09, *p*s ≥ 0.31) then declined at different rates in males and females (bin 1 > 3-6 vs bin 1 > 4-6, respectively, Bonferroni’s tests, *p*s < 0.05).

### Activated BLA-projecting DR CRF neurons co-express vGlut3 and TPH2

Experiment 3 characterized the neurochemical phenotypes of BLA-projecting CRF neurons that exhibited c-Fos expression after memory reactivation (**Fig. 4**). These neurons were selectively labeled with GFP using a CRFp-iCre-dependent approach (**Fig. 4A**). Brain tissue was collected 2 hours after a memory-reactivation session (**Fig 4A-B**). Total GFP expression in the DR did not vary by sex or memory reactivation condition (**Fig. 4C-E;** 2 x 2 ANOVA; *F*s ≤ 3.46 all *p*s ≥ 0.09), and GFP expression colocalized with CRF immunoreactivity in 98 +/- 0.2 % of the cell bodies. Memory reactivation approximately tripled the proportion of c-Fos^+^ GFP-expressing (BLA-projecting CRF) neurons independent of sex (**Fig. 4F Inset;** *t*_(14)_ = 2.18, *p* = 0.02), compared to no-memory reactivation. Among the five c-Fos^+^ GFP-expressing neuronal populations we identified (**Fig. 4F**), this response reflected increased c-Fos expression in GFP-expressing **vGlut3**^+^ neurons (vesicular glutamate transporter 3; 2 x 2 ANOVA; group main effect only, *F*_(1,12)_ = 5.48, *p* = 0.04, all other *F*s ≤ 0.34, *p*s ≥ 0.57) and a trend for increased c-Fos expression in GFP-expressing vGlut3^+^**TPH2**^+^ cells (tryptophan hydroxylase 2; group main effect only, *F*_(1,12)_ = 4.10, *p* = 0.07, all other *F*s ≤ 0.25, *p*s ≥ 0.63). Neither sex nor memory reactivation altered total immunofluorescence for these cell types (*F*s ≤ 2.57, *p*s ≥ 0.16). The GFP^+^vGlut3^+^ cell population was observed evenly across DR subregions, and the GFP^+^vGlut3^+^TPH2^+^ cell population was observed in the medial DR (**Fig. 5**); however, low cell densities prevented subregion-specific statistical analyses. Neither memory reactivation nor sex altered c-Fos expression in other GFP-expressing neuronal populations, including those that were vGlut3^-^TPH2^+^, TPH2^+^**TH**^+^ (tyrosine hydroxylase), or vGlut3^-^TPH2^-^TH^-^ (i.e., potentially multiple cell types with no or subthreshold labeling) (**Fig. 4F**; *F*s ≤ 1.74, *p*s ≥ 0.21).

**Figure 5.**
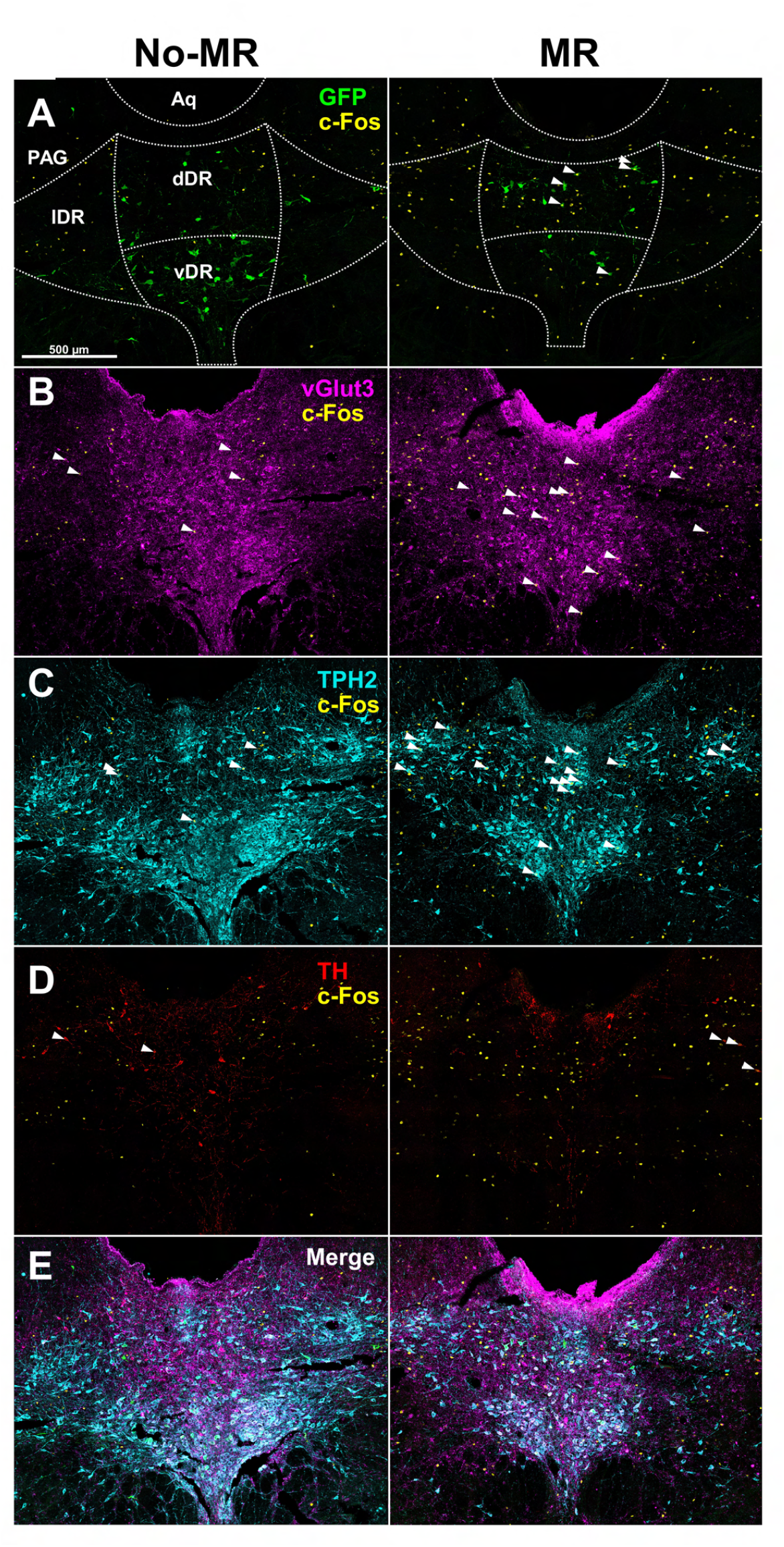
Colocalization of c-Fos with neuronal phenotype markers in the DR. Stitched confocal microscopy images of c-Fos-immunoreactivity and cell type markers in the dorsal raphe (**DR**) of representative rats in the memory reactivation (**MR**) and no-memory reactivation (**No-MR**) groups. White arrows indicate colocalization of c-Fos with a cell type marker: **(A)** GFP (BLA-projecting CRF neurons), **(B)** vGlut3 (glutamate cell marker), **(C)** TPH2 (serotonin cell marker), and **(D)** TH^+^ (dopamine cell marker in the DR). **(E)** Merged image of c-Fos and all cell type markers. **Abbreviations: Aq**, cerebral aqueduct; **dDR**, dorsal subregion of dorsal raphe; **lDR**, lateral subregion of dorsal raphe; **vDR**, ventral subregion of dorsal raphe; **PAG**, periaqueductal gray.

Memory reactivation also doubled total c-Fos expression in the DR, collapsed across GFP^+^ and GFP-*non-expressing* (i.e., non-transfected, non-BLA-projecting, and/or non-CRF) neurons, compared to no-memory reactivation (**Fig. 4G Inset**; *t*_(14)_ = 1.88, *p* = 0.02). Of the six c-Fos^+^ GFP-non-expressing neuronal populations we observed, memory reactivation produced a trend for increased c-Fos expression in the vGlut3^-^TPH2^-^TH^-^ neurons (i.e., potentially multiple cell types) independent of sex (**Fig. 4G**; group main effect only, F_(1,12)_ = 4.35, p = 0.06; all other *F*s < 2.17, *p*s > 0.17). In addition, the GFP-non-expressing TPH^+^ neuronal population exhibited greater c-Fos expression in females than males, independent of memory-reactivation condition (sex main effect only, *F*_(1,12)_ = 5.07, *p* = 0.04, all other *F*s ≤ 1.44, *p*s ≥ 0.25) despite that total TH^+^ immunofluorescence was lower in females than males (sex main effect only, *F*_(1,12)_ = 32.19, p < 0.001; all other *F*s < 0.30, *p*s > 0.60). Neither memory reactivation nor sex altered c-Fos expression in the remaining four neuronal populations (*F*s ≤ 2.82, *p*s ≥ 0.12).

## DISCUSSION

The present study investigated the involvement of the DR_CRF_ → BLA circuit in cocaine-memory maintenance. Chemogenetic DR_CRF_ → BLA circuit inhibition attenuated BLA Zif268 expression, a molecular event that is necessary for cocaine-memory reconsolidation [4,15], in the BLA, but not in the adjacent CeA (**Fig. 1**). DR_CRF_ → BLA circuit inhibition at a time point when cocaine memories were labile reduced subsequent drug-seeking behavior selectively in the cocaine-predictive context (**Fig. 2**). Conversely, DCZ treatment in the absence of hM4Di expression (**Fig. 2**) or DR_CRF_ → BLA circuit inhibition at a time point when cocaine memories were reconsolidated into long-term memory [15] (**Fig. 3**) did not alter subsequent cocaine-seeking behavior. These DCZ-induced, hM4Di-mediated, anatomically-selective, and memory-reactivation-dependent impairments in cocaine-seeking behavior indicate that DR_CRF_ → BLA circuit signaling is necessary for the maintenance of cocaine-memory strength after destabilization. BLA-projecting DR CRF neurons that were activated during cocaine-memory reconsolidation were either vGlut3^+^ or vGlut3^+^TPH2^+^ (**Fig. 4-5**), consistent with the possibility that cocaine-memory reconsolidation may involve co-release of CRF with glutamate and/or serotonin in the BLA. These findings significantly advance our understanding of the circuit-level mechanisms of cocaine-memory reconsolidation.

### DR_CRF_ → BLA in the context of brain systems involved in cocaine-memory reconsolidation

DR_CRF_ → BLA circuit inhibition likely disrupted a larger brain system that regulates cocaine-memory strength. Some elements of this system have been identified, but requisite functional connections between these elements are not well-characterized. We have demonstrated that BLA – dorsal hippocampus (**DH**) communication is necessary for cocaine-memory reconsolidation [24]. Input from the DH may facilitate protein synthesis-dependent memory re-stabilization in the BLA via a polysynaptic circuit [24], and critical input from the BLA may reach the DH through a monosynaptic circuit [29]. Another target of monosynaptic BLA projections is the nucleus accumbens (**NAc**), a brain region implicated particularly in Pavlovian drug-memory reconsolidation [5,30,31]. Chemogenetic BLA → NAc circuit inhibition disrupts Pavlovian methamphetamine-memory reconsolidation [32], and chemogenetic circuit stimulation alone (without context re-exposure) is sufficient to destabilize those memories [32]. Importantly, DR CRF likely modulates the stability of cocaine memory traces in a large network that extends well beyond the DH, BLA, and NAc. In line with this, fear memories are encoded in functionally connected microcircuits, or engrams, throughout the brain [33]. Over a hundred contextual fear engram locations have been identified in the mouse, and activation of multiple engrams corresponds to improved memory retrieval [33]. Similar network-level mechanisms are expected to regulate cocaine-memory strength, and the identification and functional characterization of underlying neural networks will be an important objective of future research.

### Cell type-specific engagement of DR in cocaine-memory reconsolidation

The DR is a neurochemically heterogenous brain region that contains CRF, dopamine, serotonin, and glutamate neurons among others [34,35]. In our study, cocaine-memory reactivation increased neuronal activation in three DR neuronal populations to varying extents (**Fig. 4F-G**). These were GFP-expressing, BLA-projecting CRF neurons and other (GFP*-non-expressing*) DR neurons that were vGlut3^+^, TPH2^+^, or vGlut3^+^TPH2^+^. These observations implicate DR CRF, glutamate, and/or serotonin release in the BLA in cocaine-memory reconsolidation, corroborating extant literature.

We have shown previously that CRF signaling in the BLA is necessary for cocaine-memory reconsolidation [16]; however, several potential sources for CRF exist in addition to the DR. In particular, CRF *diffusion* from the CeA to the BLA has been theorized to mediate the stress-induced potentiation of fear-memory consolidation [36]. CRF^+^ neurons have been reported in the BLA[37], suggesting the potential for local CRF release. Aside from the DR, a potential extra-amygdalar source of CRF is the dorsomedial prefrontal cortex (**dmPFC**). The dmPFC contains a large population of CRF neurons [27,37] and robustly projects to the BLA [38]. Furthermore, the dmPFC is involved in cocaine-memory reconsolidation [39,40]; although, its engagement appears to be paradigm-specific and may require drug exposure during memory reactivation [41]. Comprehensive understanding of the role of CRF signaling in memory reconsolidation will require the systematic consideration of multiple potential sources of CRF.

Similar to CRFR1 stimulation, glutamate receptor stimulation and synaptic plasticity in the BLA are key to cocaine-memory reconsolidation [17,42,43], but DR glutamate may not be required for this phenomenon. Multiple brain regions implicated in memory reconsolidation, including the DR, dmPFC, insular cortex, and ventral hippocampus, send glutamatergic inputs to the BLA [40,44,45]. Moreover, CRF signaling can likely stimulate local glutamate release in the BLA by exciting glutamatergic principal neuron [46,47].

Unlike CRF and glutamate, serotonin bidirectionally modulates pyramidal neuronal firing in the BLA depending on concentration and on whether the stimulation of different 5-HT receptor subtypes on principal neurons or GABAergic interneurons dominates [48,49]. Pathway-selective optogenetic stimulation of BLA-projecting serotonin neurons in the DR increases BLA neuronal firing and facilitates fear encoding via the stimulation of 5-HT1a/2a receptors [50]. Thus, increased c-Fos expression in GFP-expressing TPH2^+^ cells in the DR and subsequent serotonin release in the BLA could also support cocaine-memory reconsolidation.

The largest group of cells that exhibited a trend for increased neuronal activation during memory reconsolidation were GFP-non-expressing and vGlut3^-^TPH2^-^TH^-^ (**Fig. 4G**). These cells may be GABA neurons [51], vGlut3-negative glutamate neurons [34], or glutamate neurons with subthreshold cell type marker expression or inadequate antibody labeling (**Fig. 4G & 5**). In this initial investigation, we limited our phenotyping efforts to cell types that were expected to directly excite BLA pyramidal neurons and provided only a snapshot of neuronal activation approximately 30 minutes after memory reactivation based on c-Fos expression [28]. Future research will need to expand these analyses to additional DR cell types, additional time points to capture early and late stages of memory reconsolidation, and additional molecules relevant to cocaine-memory reconsolidation (e.g., Zif268, [4,15]).

Our report of a sex-independent mechanism for cocaine-memory reconsolidation adds to a small literature on sex-dependent [16,52] and sex-independent [16,21,53,54] mechanisms of memory reconsolidation. Females exhibited higher c-Fos expression in GFP-non-expressing TPH2^+^ DR cells (**Fig. 4G**), despite lower overall TPH2 immunoreactivity (see *Results*), relative to males, but this effect was independent of memory reactivation. Greater c-Fos expression in females might result from higher rates of serotonin neuronal firing which is modulated by sex hormones [55].

c-Fos expression in TH^+^ dopamine neurons was negligible in both sexes independent of memory reactivation. This could be due to low TH^+^ cell density in the regions of the DR we analyzed, posterior to the mesencephalic DR [56]. Low TH^+^ cell density could mask potential group differences. DR dopamine neurons encode the motivational salience of reward-predictive stimuli in the context of other information [57]. Thus, it will be particularly important to re-investigate the potential significance of TH^+^ DR neuronal populations for cocaine-memory reconsolidation in future studies.

## Conclusion

Findings in the present study conclusively demonstrate the critical contribution of DR_CRF_ → BLA circuits to cocaine-memory reconsolidation but fail to identify a single neuropeptide or neurotransmitter that controls this phenomenon. BLA-projecting DR neurons co-release CRF with serotonin and/or glutamate in the BLA. Therefore, selective chemogenetic inhibition of the DR_CRF_ → BLA circuit disrupts signaling through multiple receptor systems in the BLA. In fact, reduction in cocaine-memory strength may depend on concomitant inhibition of multiple DR neuronal populations and signaling pathways. Therapeutic inhibition of these systems may weaken abnormally strong drug memories that predispose individuals to intractable SUD [58,59]. With advancements in deep brain stimulation technology [60], our understanding of circuit-level abnormalities may be harnessed to normalize brain function and improve the lives of individuals with SUD and other neuropsychiatric disorders.

### Supplemental Methods and Materials

Supplemental methods and materials can be found on bioRxiv.

## Acknowledgements

The authors thank Robert Christian, Justine Galliou, Abigail Greenway, Peyton Krych, and Noah Frachon for expert technical assistance.

## Author Contributions

JLR: conceptualization, investigation, data analysis, visualization, writing original draft. SQ, DAS, AYP, LMA, and SKC: investigation. CWB: supervision. RAF: conceptualization, data analysis, writing original draft, review and editing, supervision, funding acquisition.

