## Supplemental Methods and Materials for "Dorsal Raphe to Basolateral Amygdala Corticotropin-Releasing Factor Circuit Regulates Cocaine-Memory Reconsolidation"

✉**Corresponding Author:** Dr. Rita A. Fuchs, Washington State University, Integrative Physiology and Neuroscience, Veterinary and Biomedical Research Building 205, PO Box 647620, Pullman, WA 99164-7620,

### Food Training

Rats received a single overnight food training session. Food-training sessions took place in standard operant-conditioning chambers (26 × 27 × 27 cm; Coulbourn Instruments, Allentown, PA) housed within sound-attenuation cubicles (Med Associates, Fairfax, Vermont, USA). Each chamber was equipped with two levers, a water bottle with sipper tube, a food tray, and a food-pellet dispenser controlled by Graphic State Notation software version 4.1.04 (Coulbourn). Water was available *ad libitum*. Responses on one lever (active lever) resulted in food reinforcement (45-mg grain-based pellets; Bio-Serv, Flemington, NJ) under a continuous reinforcement schedule. Responses on the other lever (inactive lever) had no programmed consequences. During food training, the rats had no access to the auditory or olfactory stimuli that were used during subsequent cocaine self-administration or extinction training. Training continued, as needed, until the rats obtained a minimum of 100 food pellets during a session.

### Surgery

Rats were anesthetized using ketamine and xylazine (100.0 mg/kg and 5.0 mg/kg, i.p., respectively) at least 24 hours after an overnight food training session. Back-mounted intravenous catheters were implanted into the right jugular vein as described previously [30]. Rats in Experiment 1-2 then received a unilateral virus-cocktail infusion into the median DR (30° angle, AP –7.7 mm, ML +2.88 mm, DV –5.77 mm) and stainless-steel guide cannulae implants (P1 Technologies, Roanoke, Virginia, USA) aimed bilaterally 2 mm dorsal to the BLA (AP –2.7 mm, ML +4.6, DV –6.3). The virus cocktail (200 nl) contained, in 2:1 ratio, AAV8-rCRHp-iCre (1.2 x $10^{13}$ GC/mL, Vector Biolabs, Malvern, PA) plus either AAV8-hSyn-DIO-hM4Di-mCherry (Experiment 2-3; 2.2 x $10^{13}$ GC/mL, Addgene, Watertown, MA) or AAV8-hSyn-DIO-mCherry (Control virus; 2.3 x $10^{13}$ GC/ML, Addgene) to permit the expression of hM4Di-mCherry or mCherry alone, respectively, in BLA-projecting CRF neurons. The infusion was administered over 10 minutes, and the injector was left in place for 10 minutes thereafter to minimize virus spread up the injection tract. Rats in Experiment 3 received a unilateral infusion of AAV8-rCRHp-iCre (200 nL; 1.2 x $10^{13}$ GC/mL, Vector Biolabs) into the median DR and bilateral infusions of AAVretro-hSyn-DIO-EGFP (200 nL over 2 min; 1.3 x $10^{13}$ GC/mL, Addgene) into the BLA to fluorescently label the cell bodies of BLA-projecting DR CRF neurons.

Rats received post-operative analgesic treatment for at least 48 hours after surgery and five days for post-surgical recovery. All rats received a bacon-flavored placebo tablet (5 g/tablet per os; Bio-Serv) 24 hours prior to surgery to eliminate neophobia to the postoperative analgesic. Postoperative analgesic treatment consisted of once daily oral access to a non-steroidal anti-inflammatory pain reliever (bacon-flavored Rimadyl® MD tablets containing 2 mg carprofen/5-8mg/kg). Rats received postoperative care for at least 5 days and until their weight returned to preoperative levels.

Catheters were flushed daily with an antibiotic solution of cefazolin (1.0 mg/0.1 mL; West-Ward, Eatontown, NJ; dissolved in 70 U/mL heparinized saline, Sagent Pharmaceuticals, Schaumburg, IL) followed by heparinized saline (0.1 mL, 10 U/mL) to help maintain catheter patency. The catheter and cannula were capped when not in use to prevent clogging (P1 Technologies). Catheter patency was assessed periodically using propofol (1 mg/0.1 mL; Actavis Pharma, Parsippany, NJ), a short-acting anesthetic that produces rapid, temporary loss of muscle tone when administered intravenously.

### Drug self-administration training

Drug self-administration training was conducted six days per week in operant-conditioning chambers (Coulbourn) configured to form two contexts that contained distinctly different auditory, visual, tactile, and olfactory modalities. The specific contexts used for self-administration training versus extinction training were counterbalanced across rats. One context consisted of a continuous red house light (0.4-fc brightness), intermittent pure tone (80 dB, 1 kHz; 2 sec on, 2 sec off), pine scented air freshener (Car Freshener Corp., Watertown, NY), and wire-mesh flooring (26 cm × 27 cm). The other context consisted of an intermittent white stimulus light over the inactive lever (1.2-fc brightness; 2 sec on, 2 secs off), continuous pure tone (75 dB, 2.5 kHz), vanilla-scented air freshener (Sopus Products, Moorpark, CA), and a slanted ceramic tile wall that bisected the bar flooring (19 cm × 27 cm).

Cocaine self-administration training sessions took place 2 hours per day during the rats' dark phase. Active-lever responses resulted in cocaine reinforcement (0.5 mg/kg per 50-µL infusion, delivered i.v. over 2 sec; generously provided by the NIDA Drug Supply Program, Research Triangle Park, NC) under a fixed ratio 1 schedule with a 20-sec timeout period after each infusion. Inactive-lever responses were not reinforced at any time. Training continued until rats received a minimum of 10 sessions during which they obtained ≥ 10 infusions/session.

### Extinction training

Rats received seven daily two-hour extinction training sessions in the other context. During the extinction training sessions, lever responses were not reinforced. The number of extinction sessions were set to seven to match memory age at the time of memory reactivation across rats. Immediately after extinction session 4, the rats in Experiments 1-2 were acclimated to the intracranial infusion procedure. The stainless-steel injection cannulae (33-gauge; P1 Technologies) were inserted into the guide cannula to a depth of 2 mm past the tip of the guide cannula. The injection cannulae remained in place for 4 minutes with no infusion of fluids while the rats were held gently by the experimenter.

#### Memory reactivation

Rats were placed into the cocaine-predictive context for 15 minutes to elicit the retrieval and destabilization of contextual cocaine memories (memory reactivation) or remained in their home cages (no-memory reactivation). Assignments to these experimental conditions were balanced based on active-lever responding during the last three cocaine self-administration sessions. The 15-minute session length is sufficient to destabilize cocaine memories without eliciting overt behavioral extinction [1,2]. Rats in the memory reactivation groups were connected to the infusion lines, but lever responses were not reinforced. Rats received intra-BLA infusions of vehicle (Saline; 0.5 µl/hemisphere) or DCZ (0.1 mM; HB9126, Hello Bio, Princeton, NJ) administered either immediately after or six hours after the memory-reactivation session (Experiments 1 and 2, respectively), or they were euthanized two hours after the memory-reactivation session without intracranial treatment (Experiment 3).

#### Tests of memory strength

Starting one day after memory reactivation, rats in Experiments 1-2 received daily two-hour extinction training sessions until their active-lever responding declined to $\leq$ 25 on two consecutive days. They were then placed into the cocaine-predictive context for two hours. The rats were attached to the infusion lines in the cocaine-predictive context. Lever responses were not reinforced in either context. Active-lever responses in the extinction and in the cocaine-predictive contexts were used as indices of extinction and cocaine-memory strength, respectively.

#### DREADD functional validation

At least 3 days after testing in the cocaine-predictive context, a subset of rats from Experiments 1-2 received a memory-reactivation session followed immediately by intra-BLA VEH or DCZ treatment. Rats were perfused two hours later, and brain tissue was collected to assess Zif268 expression. Assignment to treatment conditions was counter balanced to avoid repeated DCZ treatment.

#### Brain histology

Rats received an overdose of ketamine hydrochloride and xylazine (100 and 5 mg/kg, i.v., or 300 and 15 mg/kg, i.p., respectively) and were then transcardially perfused first with ice-cold phosphate buffered saline (PBS, 200 mL) then with 4% paraformaldehyde dissolved in PBS (PFA, 200 mL). The brains were post-fixed overnight in PFA at 4 °C. The brains were extracted, cryoprotected in 30% sucrose solution containing 0.1% sodium azide, and stored at −80 °C. Brains were sectioned on a cryostat (Leica Biosystems, Buffalo Grove, Illinois, USA). Thirty-µm brain sections were stained with DAPI (1:1000; D1306, Invitrogen) and mounted on glass slides. Injection-cannula placement and virus spread were assessed using an epifluorescence microscope (Leica CTR 6500, Wetzler, Germany).

#### *In situ* hybridization

To examine the specificity of the CRF antibody (**Fig. 1B**), we examined the extent to which CRF mRNA expression colocalized with CRF immunoreactivity (1067 cells counted; *n* = 2 males). For *in situ* hybridization, three digoxigenin-labeled mRNA probes (MicroSynth, Balgach, Switzerland) were designed to be directed against different regions of rat CRF mRNA (NCBI GenBank reference sequence: NM_031019.2). The selected probes were predicted to 1) be unfolded (using RNA Webserver), 2) have melting temperatures between 65-75°C, 3) be comprised of 40 to 60% guanine and cytosine, and 4) have lower than 0.001 affinity for non-CRF mRNA (NCBI BLAST). The three RNA probes best meeting these criteria were: 1) 5'-CAA CCU CAG CCG AUU CUG AU-3', 2) 5'-CGG CUA ACU UUU UCC GCG UG-3', and 3) 5'-GAG AGC CUA UAU ACC CCU UA-3'. The brain slices were serially washed with: 1) 0.2 M diethyl pyrocarbonate (DEPC)-treated PBS, 2) 3% peroxide; 3) 0.2 M hydrochloric acid, and 4) 0.25% acetic anhydride. The mRNA probes (0.1 µg/mL) were hybridized with CRF mRNA at 60°C in a solution containing hybridization buffer [5x standard citrate saline (SSC), pH 7.0; 50 µg/mL heparin; 0.001 M EDTA, pH 8.0; 50% molecular grade formamide, deionized, 1x Denhart's solution, 0.1% Tween-20, 0.25 mg/mL yeast tRNA, and 10% dextran sulphate, brought to final volume in DEPC-treated water] for 20 hours. Following probe hybridization, the brain slices were washed with gradients of SSC, and unbound probes were digested with RNAse A (20 µg/mL; Roche, Indianapolis, IN). Next, the brain slices were incubated with sheep anti-digoxigenin-POD (1:100; 11207733910, Roche, Basel, Switzerland) with agitation overnight at room temperature. Signal was then amplified by TSA Plus cyanine 3 (NEL744001KT; Akoya Biosciences, Marlborough, MA). Brain slices were washed in PBS (4 x 3 minutes) and placed in blocking solution containing 5% normal donkey serum and 0.1% Triton X-100 for 45 minutes.

#### Immunohistochemistry

Immunohistochemistry assays are described below in the order they appear in the manuscript. In each experiment, brains from transcardially perfused rats were sectioned in the coronal plane at 30-µm thickness. The brain slices were mounted on charged glass slides and cover slipped with Prolong Diamond Antifade (P36965, Thermo Fisher) after immunolabeling. The edges of the coverslips were sealed with clear nail polish, and the slides were stored at 4 °C.

***Validation of the CRF antibody.*** To evaluate colocalization with CRF immunoreactivity (**Fig. 1A**), the brain slices were then incubated with guinea pig anti-CRF polyclonal anti-serum (1:1000; T-5007, BMA Biomedicals, Augst, Switzerland) at 4 °C with agitation overnight. The brain slices were washed in PBS (4 x 3 minutes), then incubated with secondary antibody solution containing goat anti-guinea pig 568 IgG (1:200; A11075, Thermo Fisher) for 1.5 hours at room temperature, followed by a final wash in PBS (4 x 3 min).

***Validation of the cell specific- and pathway-specific viral approach.*** To examine the selectivity of the viral approach (**Fig. 1B-E**), we evaluated the extent to which CRFp-iCRE-dependent hM4Di-mCherry or mCherry expression colocalized with CRF immunoreactivity in the DR. Brain slices were placed in blocking solution containing 5% normal goat serum and 0.5% Triton X-100 in 0.5% PBS (PBST) for 1 hour. They were then incubated with primary antibody solution that contained guinea pig anti-CRF polyclonal anti-serum (1:700; T-5007) in the blocking solution at 4 °C for 72 hours. The brain slices were then washed in PBS (4 x 3 minutes), then incubated with secondary antibody solution that containing goat anti-guinea pig 647 polyclonal IgG (1:250; 106-605-003, Jackson ImmunoResearch) for two hours at room temperature, followed by a final wash in PBS (4 x 3 min).

***Validation of DREADD functionality.*** To examine whether DCZ was sufficient to stimulate functionally active hM4Di (**Fig. 1F-H**), changes in Zif268 expression were examined in rats expressing hM4Di-mCherry or mCherry alone two hours after a 15-minute memory reactivation session followed by intra-BLA treatment with DCZ (0.1 mM) or vehicle (Saline; 0.5 µl/hemisphere). Brain slices were placed in blocking solution for one hour. They were then incubated with primary antibody solution containing rabbit anti-EGR1 (i.e., Zif268) monoclonal IgG (1:1000; 15F7, Cell Signaling Technology, Danvers, MA) and chicken anti-mCherry polyclonal IgY (1:1000; MCHERRY-0100, Avēs Labs, Davis, CA) at room temperature for 24 hours. The brain slices were then washed in PBS (4 x 3 minutes), then incubated in secondary antibody solution containing goat anti-rabbit 488 polyclonal IgG (1:250; 111-545-003, Jackson ImmunoResearch) and goat anti-chicken 647 polyclonal IgY (1:250; 103-605-155, Jackson ImmunoResearch) at room temperature for 2 hours, followed by a final wash in PBS.

***DR c-Fos immunolabeling and cell phenotyping**.* To determine the neurochemical phenotype of cFos-expressing DR cells (**Fig. 4-5**), immunolabeling of multiple targets were completed sequentially over 3 days to avoid antibody interactions. *On day 1,* brain slices were placed in blocking solution for one hour. They were then incubated with primary antibody solution containing sheep anti-TH polyclonal IgG (1:1000; AB1542, Sigma-Aldrich, St Louis, MO) at room temperature for 24 hours. *On day 2,* the brain slices were washed in PBS (4 x 3 minutes) then incubated with secondary antibody solution containing donkey anti-sheep 594 polyclonal IgG (1:250; 713-585-147, Jackson ImmunoResearch) in PBS at room temperature for two hours. They were then washed in PBS (4 x 3 minutes) and placed in blocking solution containing 5% normal goat serum and 0.5% PBST for 1 hour. They were then incubated with primary antibody solution containing guinea pig anti-vGlut3 polyclonal IgG (1:500; Synaptic Systems, Göttingen, Germany) in blocking solution, at room temperature for 24 hours. *On day 3,* the brain slices were washed in PBS (4 x 3 minutes) then incubated with secondary antibody solution containing goat anti-guinea pig 647 polyclonal IgG (1:250; 106-605-003, Jackson ImmunoResearch) in PBS at room temperature for two hours. They were then washed in PBS (4 x 3 minutes) and then incubated with primary antibody solution containing chicken anti-c-Fos monoclonal IgY (1:500; 226009, Synaptic Systems), rabbit anti-TPH2 polyclonal IgG (1:1000; NB100-74555, Novus Biologicals, Centennial, CO), and goat anti-GFP polyclonal IgG conjugated to Dylight™ 488 (1:1000; 600-141-215, Rockland, Limerick, PA) in PBS at room temperature for 24 hours. *On day 4*, the brain slices were washed in PBS (4 x 3 minutes) and then incubated with secondary antibody solution containing goat anti-chicken Cy3 polyclonal IgY (1:250; 103-165-155, Jackson ImmunoResearch) and goat anti-rabbit 405 polyclonal IgG (1:250; 111-475-003, Jackson ImmunoResearch) in PBS at room temperature for two hours, followed by a final wash in PBS.

**Image Analysis**

The slides were imaged using an epifluorescence microscope (10x dry; Leica CTR 6500) or confocal microscopes (40x or 63x oil immersion; Leica SP8; 20x or 63x; Nikon A1R+, Melville, NY). For Experiment 1, single-labeled cell nuclei were quantified on four images per subject using a custom macro in ImageJ. The threshold for counting a cell was calibrated to include only bright nuclear labeling. For Experiment 3, multi-label cell bodies were counted on three z-stacks per DR subregion per subject. Each z-stack spanned 28 µm with 1 µm steps. Multi-labeled cell bodies were quantified in ImageJ by two independent observers blinded to the subject's treatment condition. The extent of co-localization between c-Fos-immunoreactivity and GFP expression (BLA-projecting CRF neuronal label), and vesicular glutamate transporter 3 (vGlut3; glutamatergic neuronal marker), tryptophan hydroxylase 2 (TPH2; 5-HT neuronal marker), and tyrosine hydroxylase (TH; dopaminergic neuronal marker in the DR [3]) immunoreactivity was assessed in the DR. Only cells that had clear nuclear labeling were counted as c-Fos-positive. Inter-observer reliability was high ($R^2$ = 0.96). All data were converted to density values (cell bodies/mm2) and averaged across all replicates

| Table 1S. Virus constructs, antibodies, and staining used in Experiments 1-3 | | | | |
|---|---|---|---|---|
| **Type** | **Designation** | **Manufacturer/ Supplier** | **Product Identifier/ RRID** | **Concentration/ Dilution** |
| Virus | AAV8-rCRHp-iCre | Vector Biolabs | Service ID: 7600 | 1.2 x $10^{13}$ GC/mL |
| Virus | AAV8-hSyn-DIO-hM4Di-mCherry | Addgene | Plasmid #44362 Addgene_44362 | 2.2 x $10^{13}$ GC/mL |
| Virus | AAV8-hSyn-DIO-mCherry | Addgene | Plasmid #50459 Addgene_50459 | 2.3 x $10^{13}$ GC/mL |
| Virus | AAVretro-hSyn-DIO-EGFP | Addgene | Plasmid #50457 Addgene_50457 | 1.3 x $10^{13}$ GC/mL |
| Primary Antibody | sheep anti-Digoxigenin-POD | Roche | 11207733910 AB_514500 | 1:100 |
| Primary Antibody | mouse anti-c-Fos monoclonal IgG | Santa Cruz Biotechnology | sc-271243 AB_10610067 | 1:500 |
| Primary Antibody | guinea pig anti-CRF polyclonal | BMA Biomedicals | T-5007 AB_518256 | 1:700 |
| Primary Antibody | rabbit anti-EGR1 monoclonal IgG | Cell Signaling Technology | 15F7 AB_2097038 | 1:1000 |
| Primary Antibody | chicken anti-mCherry polyclonal IgY | Avēs Labs | MCHERRY-0100 AB_2910557 | 1:1000 |
| Primary Antibody | sheep anti-TH polyclonal IgG | Sigma-Aldrich | AB1542 AB_90755 | 1:1000 |
| Primary Antibody | guinea pig anti-vGlut3 polyclonal IgG | Synaptic Systems | 135 204 AB_2619825 | 1:500 |
| Primary Antibody | chicken anti-c-Fos monoclonal IgY | Synaptic Systems | 226009 AB_2943525 | 1:500 |
| Primary Antibody | rabbit anti-TPH2 polyclonal IgG | Novus Biologicals | NB100-74555 AB_1049988 | 1:1000 |
| Primary Antibody | goat anti-GFP polyclonal IgG conjugated to Dylight™ 488 | Rockland | 600-141-215 AB_1961516 | 1:1000 |
| Secondary Antibody | goat anti-mouse 647 polyclonal IgG | Jackson ImmunoResearch | 115-605-003 AB_2338902 | 1:250 |
| Secondary Antibody | goat anti-guinea pig 488 polyclonal IgG | Invitrogen | A-11073 AB_2534117 | 1:250 |
| Secondary Antibody | goat anti-guinea pig 647 polyclonal IgG | Jackson ImmunoResearch | 106-605-003 AB_2337446 | 1:250 |
| Secondary Antibody | goat anti-rabbit 488 polyclonal IgG | Jackson ImmunoResearch | 111-545-003 AB_2338046 | 1:250 |
| Secondary Antibody | goat anti-chicken 647 polyclonal IgY | Jackson ImmunoResearch | 103-605-155 AB_2337392 | 1:250 |
| Secondary Antibody | donkey anti-sheep 594 polyclonal IgG | Jackson ImmunoResearch | 713-585-147 AB_2340748 | 1:250 |
| Secondary Antibody | goat anti-chicken Cy3 polyclonal IgY | Jackson ImmunoResearch | 103-165-155 AB_2337386 | 1:250 |
| Secondary Antibody | goat anti-rabbit 405 polyclonal IgG | Jackson ImmunoResearch | 111-475-003 AB_2338035 | 1:250 |
| Nuclear Stain | DAPI | Invitrogen | D1306 | 1:1000 |

### Table S2. Behavioral history statistics for Experiments 1-3.

| Experiment 1 | Self-administration[a] | | | | | | | | | Extinction[b] | | | | | | Memory Reactivation[c] | | | | | | Post Hoc Tests[d] |
|---|---|---|---|---|---|---|---|---|---|---|---|---|---|---|---|---|---|---|---|---|---|---|
| Dependent Measures | Active Lever | | | Inactive Lever | | | Cocaine Infusions | | | Active Lever | | | Inactive Lever | | | Active Lever | | | Inactive Lever | | | |
| | *df* | *F* | *p* | *df* | *F* | *p* | *df* | *F* | *p* | *df* | *F* | *p* | *df* | *F* | *p* | *df* | *F* | *p* | *df* | *F* | *p* | **Self-administration**: Inactive d1 > d4-7,9,10; Infusions d1 < d8-10. |
| **Sex** | 1,48 | 1.03 | 0.32 | 1,48 | 2.68 | 0.11 | 1,48 | 2.57 | 0.12 | 1,49 | 0.71 | 0.40 | 1,49 | 2.24 | 0.14 | 1,49 | 1.26 | 0.27 | 1,49 | 4.14 | 0.05 | |
| **Group** | 2,48 | 0.13 | 0.88 | 2,48 | 1.28 | 0.29 | 2,48 | 1.04 | 0.36 | 2,49 | 1.68 | 0.20 | 2,49 | 0.94 | 0.40 | 2,49 | 0.38 | 0.69 | 2,49 | 0.81 | 0.45 | **Extinction:** |
| **Day** | 9,432 | 3.12 | <0.001 | 9,432 | 6.30 | <0.001 | 9,432 | 22.87 | <0.001 | 6,294 | 42.23 | <0.001 | 6,294 | 26.45 | <0.001 | | | | | | | Active d1 > d2-7; |
| **Sex x group** | 2,48 | 0.12 | 0.89 | 2,48 | 0.84 | 0.65 | 2,48 | 0.14 | 0.87 | 2,49 | 1.45 | 0.24 | 2,49 | 0.46 | 0.63 | 2,49 | 2.22 | 0.12 | 2,49 | 2.04 | 0.14 | male inactive d1 > d2-7. |
| **Sex x day** | 9,432 | 1.09 | 0.37 | 9,432 | 0.48 | 0.62 | 9,432 | 0.34 | 0.96 | 6,294 | 0.11 | 1.00 | 6,294 | 2.68 | 0.02 | | | | | | | **Memory Reactivation:** |
| **Group x day** | 18,432 | 1.11 | 0.34 | 18,432 | 0.39 | 0.94 | 18,432 | 1.50 | 0.09 | 12,294 | 0.91 | 0.54 | 12,294 | 1.05 | 0.40 | | | | | | | Inactive: male < female |
| **Sex x group x day** | 18,432 | 0.74 | 0.77 | 18,432 | 1.07 | 0.38 | 18,432 | 0.99 | 0.47 | 12,294 | 1.61 | 0.09 | 12,294 | 1.66 | 0.07 | | | | | | | |

| Experiment 2 | Self-administration[a] | | | | | | | | | Extinction[b] | | | | | | Memory Reactivation[c] | | | | | | Post Hoc Tests[d] |
|---|---|---|---|---|---|---|---|---|---|---|---|---|---|---|---|---|---|---|---|---|---|---|
| Dependent Measures | Active Lever | | | Inactive Lever | | | Cocaine Infusions | | | Active Lever | | | Inactive Lever | | | Active Lever | | | Inactive Lever | | | **Self-administration:** |
| | *df* | *F* | *p* | *df* | *F* | *p* | *df* | *F* | *p* | *df* | *F* | *p* | *df* | *F* | *p* | *df* | *F* | *p* | *df* | *F* | *p* | Infusions d1 < d10. |
| **Sex** | 1,16 | 0.80 | 0.39 | 1,16 | 0.17 | 0.69 | 1,16 | 1.77 | 0.20 | 1,17 | 0.01 | 0.93 | 1,17 | 3.71 | 0.07 | 1,17 | 1.26 | 0.28 | 1,17 | 0.00 | 0.98 | **Extinction:** |
| **Group** | 1,16 | 1.71 | 0.21 | 1,16 | 0.79 | 0.39 | 1,16 | 0.38 | 0.55 | 1,17 | 1.98 | 0.18 | 1,17 | 0.48 | 0.50 | 1,17 | 4.25 | 0.06 | 1,17 | 0.02 | 0.89 | Active d1 > d2-7 |
| **Day** | 9,144 | 3.60 | <0.001 | 9,144 | 2.36 | 0.02 | 9,144 | 4.24 | <0.001 | 6,102 | 21.37 | <0.001 | 6,102 | 3.84 | 0.002 | | | | | | | |
| **Sex x group** | 1,16 | 0.61 | 0.45 | 1,16 | 0.14 | 0.71 | 1,16 | 1.14 | 0.30 | 1,17 | 0.01 | 0.93 | 1,17 | 3.11 | 0.10 | 1,17 | 2.45 | 0.14 | 1,17 | 0.01 | 0.94 | |
| **Sex x day** | 9,144 | 1.16 | 0.32 | 9,144 | 1.29 | 0.25 | 9,144 | 0.74 | 0.67 | 6,102 | 0.53 | 0.79 | 6,102 | 0.63 | 0.70 | | | | | | | |
| **Group x day** | 9,144 | 1.40 | 0.20 | 9,144 | 1.16 | 0.33 | 9,144 | 0.70 | 0.71 | 6,102 | 0.86 | 0.52 | 6,102 | 0.32 | 0.93 | | | | | | | |
| **Sex x group x day** | 9,144 | 0.84 | 0.58 | 9,144 | 1.66 | 0.10 | 9,144 | 0.53 | 0.85 | 6,102 | 0.09 | 1.00 | 6,102 | 1.31 | 0.26 | | | | | | | |

| Experiment 3 | Self-administration[a] | | | | | | | | | Extinction[b] | | | | | | Memory Reactivation[c] | | | | | | Post Hocs Tests[d] |
|---|---|---|---|---|---|---|---|---|---|---|---|---|---|---|---|---|---|---|---|---|---|---|
| Dependent Measures | Active Lever | | | Inactive Lever | | | Cocaine Infusions | | | Active Lever | | | Inactive Lever | | | Active Lever | | | Inactive Lever | | | **Self-administration:** |
| | *df* | *F* | *p* | *df* | *F* | *p* | *df* | *F* | *p* | *df* | *F* | *p* | *df* | *F* | *p* | *df* | *t* | *p* | *df* | *t* | *p* | Infusions d2 < d10 |
| **Sex** | 1,12 | 2.46 | 0.14 | 1,12 | 0.97 | 0.35 | 1,12 | 2.57 | 0.14 | 1,13 | 1.48 | 0.25 | 1,13 | 0.45 | 0.52 | 7 | 1.31 | 0.23 | 7 | 0.08 | 0.94 | **Extinction:** |
| **Group** | 1,12 | 3.29 | 0.10 | 1,12 | 0.80 | 0.39 | 1,12 | 1.08 | 0.32 | 1,13 | 1.49 | 0.24 | 1,13 | 0.87 | 0.37 | | | | | | | Active d1 < d2-7 |
| **Day** | 9,108 | 1.49 | 0.16 | 9,108 | 3.42 | <0.001 | 9,108 | 5.36 | <0.001 | 6,78 | 24.27 | <0.001 | 6,78 | 8.03 | <0.001 | | | | | | | |
| **Sex x group** | 1,12 | 3.13 | 0.10 | 1,12 | 0.15 | 0.70 | 1,12 | 0.20 | 0.66 | 1,13 | 0.70 | 0.42 | 1,13 | 0.01 | 0.94 | | | | | | | |
| **Sex x day** | 9,108 | 0.83 | 0.44 | 9,108 | 0.37 | 0.95 | 9,108 | 1.00 | 0.44 | 6,78 | 0.17 | 0.99 | 6,78 | 0.42 | 0.87 | | | | | | | |
| **Group x day** | 9,108 | 0.98 | 0.47 | 9,108 | 0.23 | 0.46 | 9,108 | 0.56 | 0.82 | 6,78 | 0.75 | 0.61 | 6,78 | 1.71 | 0.13 | | | | | | | |
| **Sex x group x day** | 9,108 | 1.18 | 0.31 | 9,108 | 1.15 | 0.33 | 9,108 | 0.76 | 0.66 | 6,78 | 1.94 | 0.08 | 6,78 | 0.18 | 0.98 | | | | | | | |

[a]Active- and inactive-lever responses and cocaine infusions across the last 10 self-administration training days were analyzed using 2 x 3 x 10 or 2 x 2 x 10 mixed-factorial ANOVAs, where appropriate. [b]Active- and inactive-lever responses across the seven extinction training days were analyzed using 2 x 3 x 7 or 2 x 2 x 7 mixed-factorial ANOVAs, where appropriate. [c]The total number of lever responses during the 15-min memory-reactivation session was analyzed using 2 x 3 ANOVAs, 2 x 2 ANOVAs, or independent samples t-tests, where appropriate. [d]Significant Bonferroni post hoc tests.

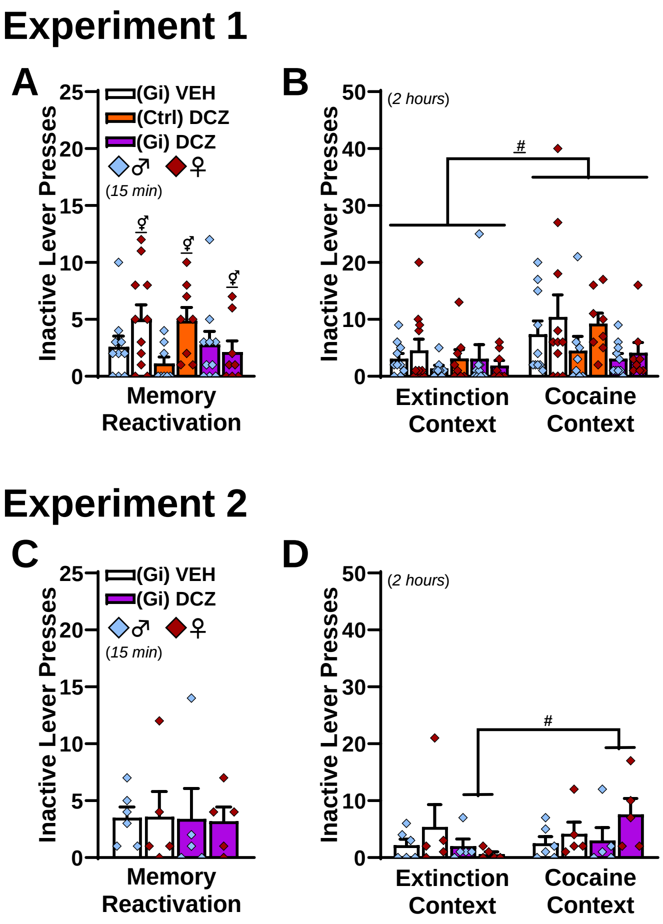


**Supplemental Figure 1. Inactive-lever responding during memory reactivation and at test in Experiments 1-2.** Inactive-lever responding remained low in both experiments. **(A)** Inactive-lever responding (mean ± SEM) during the 15-minute memory-reactivation session in Experiment 1. There were no pre-existing differences between the subsequent treatment groups in inactive-lever responding, but females responded more than males (2 x 3 ANOVA, ⚥sex main effect only, $F_{(1,49)} = 4.14$, $p = 0.05$; all other $F$s $\leq 2.04$, $p$s $\geq 0.14$). **(B)** Inactive-lever responding (mean ± SEM) during the 2-hour test sessions in the extinction and cocaine-predictive contexts in Experiment 1. The cocaine-predictive context elicited more inactive-lever responding than the extinction context, independent of sex or treatment group (2 x 3 x 2 ANOVA, #context main effect only, $F_{(1,49)} = 10.88$, $p = 0.002$; all other $F$s $< 2.05$, $p$s $> 0.14$). **(C)** Inactive-lever responding (mean ± SEM) during the 15-min memory-reactivation session in Experiment 2. Inactive-lever responding did not differ as a function of sex or subsequent treatment group (2 x 2 ANOVA, all $F$s $< 0.02$, all $p$s $> 0.89$). **(D)** Inactive-lever responding (mean ± SEM) at test in the extinction and cocaine-predictive contexts in Experiment 2. The cocaine-predictive context elicited more inactive-lever responding than the extinction context following delayed DCZ treatment, independent of sex (2 x 2 x 2 ANOVA, treatment x context interaction effect, $F_{(1,17)} = 5.19$, $p = 0.04$, #Bonferroni's test, $p < 0.05$; all other $F$s $< 3.74$, $p$s $> 0.07$).
